# Environment and plant genetics shape barley rhizosphere microbiome structure across contrasting locations

**DOI:** 10.64898/2026.07.20.739634

**Authors:** Erik Z. Killian, Jessica L. Williams, Anna Halpin-McCormick, Patrick M. Ewing, Michael B. Kantar, Jennifer Lachowiec, Jamie D. Sherman, Jed O. Eberly

## Abstract

Soil microorganisms are crucial for plant survival and productivity, but factors governing rhizosphere recruitment across diverse regions remain unclear. This study investigated the rhizosphere microbiome of barley, using elite cultivars across seven location-year trials to evaluate the effects of environmental factors and crop genotype on bacterial and fungal community composition. Three locations were in the US northern Great Plains, and Hawai’i was used as a contrasting environment. A greenhouse reciprocal transplant study determined the relative contributions of soil physicochemical factors and soil inoculum to rhizosphere community structure. Using 16S and ITS2 amplicon sequencing, the study characterized bacterial and fungal microbiomes and assessed the contribution of environment, soil chemistry, and barley genetics to microbial community assembly. In locations within the adapted range of barley, Actinobacteriota was the dominant phylum, while Proteobacteria was dominant in Hawai’i. Variance partitioning showed that 73% of bacterial and 80% of fungal genera were associated with location-year effects while 53% of bacterial and 36% of fungal genera were responsive to soil factors. Enrichment analysis found 21.6% of bacterial and 51.4% of fungal ASVs were unique to specific barley genetic subpopulations. Results from the reciprocal transplant study validated field observations by demonstrating that 20.7% of the variation in community structure was explained by soil while 18.2% was explained by inoculum source. These findings demonstrate that environmental variation is the dominant constraint on rhizosphere community composition but within these constraints, barley genotype drives recruitment of distinct bacterial and fungal taxa.

**Importance:** These findings underscore the complex interplay between plant genotype, environment, and microbial community assembly, providing insights into how barley recruits distinct microbial communities in the rhizosphere across different environments. These insights have the potential to be leveraged for management and plant breeding strategies to optimize plant-microbe interactions for enhancing agricultural sustainability.

## 1. Introduction

Soil microorganisms are integral to plant productivity, contributing to nutrient cycling and stress protection (Liu et al., 2023a; Wall et al., 2015). Soil hosts plant-associated microbes, including bacteria and fungi, which interact directly and indirectly with plant roots (Chepsergon and Moleleki, 2023). These microbes enhance crop performance by improving nutrient availability and suppressing plant-pathogens (Liu et al., 2021b; Zhao et al., 2021), and thus influence the fitness of plant hosts within a given environment. Despite these impacts, the interplay of stochasticity, environmental factors, and plant genetics in shaping communities and crop productivity and sustainability remain poorly understood.

The ability of a plant to recruit microbes is limited by the available soil community, which is shaped by the environment (Coller et al., 2019). Global soil surveys have revealed broad patterns of microbial composition with communities generally consisting of a few dominant groups and many rarer groups, and plants selectively recruit a subset of soil microbes to the rhizosphere, creating a distinct and less diverse community in the rhizosphere (Santoyo, 2022; Walker et al., 2003). Plant genetics govern this recruitment through the production of root exudates, which induce community-level changes to filter and select specific taxa (Agler et al., 2016; Banerjee et al., 2018; Berry and Widder, 2014; Bulgarelli et al., 2013). Species-specific differences in rhizosphere community recruitment have been observed, even at relatively narrow plant phylogenetic scales (Erlandson et al., 2018; Grayston et al., 1998). Recent studies have identified specific plant genes impacting the rhizosphere microbiome through molecular signaling and root exudate production (Liu et al., 2023b), and studies in rice, maize, sorghum, and switchgrass have identified plant genetic loci associated with recruitment of beneficial microbes (Deng et al., 2021; Edwards et al., 2023; Yu et al., 2021; Zhang et al., 2019; Zhang et al., 2025).

Crop genotypes are shaped by breeding programs that prioritize traits such as yield, grain quality, and disease and herbicide resistance (Smith et al., 2020). While breeding programs favor broad adaptation, genotype performance often depends on local temperature, photoperiod, soil fertility, and other environmental constraints, so varieties can respond differently when grown outside this adapted range (Ewing et al., 2023; Lawn et al., 1995; Teixeira et al., 2015). Breeding programs focus on target production environments to ensure crop productivity within a range of environmental conditions which has resulted in widely adapted genotypes, but this adaptation is often limited to regions with similar climate conditions, soil fertility, and yield potential (Ceccarelli, 1989; Smith et al., 2020).

Current evidence suggests that rhizosphere assembly reflects both deterministic processes, such as environmental and host genetics, and stochastic processes, but their relative importance remains uncertain (Dini-Andreote et al., 2015; Ma et al., 2025; Mo et al., 2024; Rain-Franco et al., 2024; Xie et al., 2024; Ye et al., 2024). This distinction is important because deterministic processes may structure dominant, generalist taxa, while stochasticity may be more important for rare, specialist taxa that may have large impacts on plant fitness (Li et al., 2025; Riddley et al., 2025).

To gain insight into the role these processes play in shaping microbial communities, we performed a common garden experiment with a population of 232 barley (*Hordeum vulgare* L.) genotypes grown across contrasting soil and climatic environments, including a location outside the adapted range of the barley genotypes. We hypothesized that i) environmental conditions would be the dominant driver of barley rhizosphere microbiome structure while plant genotype would primarily influence specific taxa rather than overall community composition; ii) growth outside the adapted range would increase environmental filtering, resulting in rhizosphere communities that diverge from those in adapted environments; and iii) soil physicochemical properties and resident soil microbiota would make distinct contributions to rhizosphere assembly. The objectives of this study were to i) characterize rhizosphere bacterial and fungal community variation across a large barley population within and outside adapted environments and (ii) experimentally determine the relative contributions of soil physicochemical factors and resident soil microbiota to rhizosphere community assembly through a reciprocal transplant study.

## 2. Materials and Methods

### 2.1. Site characteristics and field experiments

Common garden field studies were performed in 2021 and 2022 at the Post Research Farm, Bozeman, MT; the Central Agricultural Research Station (CARC), Moccasin, MT; the Eastern South Dakota Soil and Water Research Farm, Brookings, SD; and in 2021, at the Hawaii Agriculture Research Center in Kunia, Oahu, HI. At each field site, 232 elite barley lines from the S2MET population (Neyhart et al., 2019) were randomly assigned to 12 blocks in an augmented block design. The lines in this population are representative of the phenotypic and genetic diversity in North American barley breeding programs. They are descended from elite parents chosen from four public and one private breeding program that target the US northern Great Plains. Four common grown Montana barley varieties, Hockett, Odyssey, Merit-57, and Lavina, were replicated once per block as controls. Locations in MT and SD represent barley growing regions of the Northern Great Plains and are representative of environments included in barley breeding programs in this region. Hawaii, with its distinct soils, climate, and diurnal patterns, was selected as a contrasting environment far outside the adapted environments represented in the barley breeding programs. The lack of adaptation was confirmed by the failure of most barley lines to flower due to photoperiod sensitivity (data not shown). At each field site, location specific management practices for fertility and weed control were followed and neighboring fields were used between years. At the Post Farm (swMT), was planted in three-row, 3.7 m plots with 30.5 cm row spacing, following a fallow-oats (*Avena sativa* L.)-barley rotation. At CARC (cntrMT) plots were 5 rows, 4.9 m long with 30.5 cm row spacing. Field history at cntrMT varied by growing year, where the 2021 field had mustard (*Brassica napus* L.) followed by oats and the 2022 field had previous crops of oats followed by mustard. The Brookings, SD location was planted in 7-row, 4.6 m long plots with 30.5 cm row spacing, following a soybean (*Glycine max* L.) crop. At Kunia, Oahu, HI, the field history was 3 years of papaya (*Carica papaya* L.) followed by 1 year of fallow. Climate and soil physical and chemical properties are shown in Table S1. Plots were harvested by combine after grain ripening, and seed was weighed for gross yield calculations. Test weight, percent plump grain, and percent grain protein were measured on cleaned grain samples from each plot.

### 2.2. Sample collection and processing

Rhizosphere sampling was performed when plants reached stem elongation phase. In each plot, six plants were dug up with a trowel to a depth of 15 cm while maintaining as much of the intact root system as possible. Above ground plant biomass was separated and retained for biomass measurements. After lightly shaking off loose soil, roots were placed in ZipLock bags and stored on ice in coolers for transport. Soil samples (0-15 cm) were collected at the same for bulk soil microbial community analysis. A subsample was transported on ice and stored at -70°C for DNA extractions. Additional samples were collected, dried, sieved and submitted to Ward Laboratories Inc. (Kearny, NE) for chemical analysis.

Roots were sonicated for one minute in a 0.9% NaCl solution to remove the rhizosphere soil. The solution was passed through a 600 µm sieve to remove rocks and root tissues and centrifuged at 2000xg for 2 minutes, and the resulting pellet of rhizosphere soil was stored at - 70° C. DNA extractions were performed on rhizosphere soil using the Qiagen DNeasy PowerSoil Pro DNA Isolation Kit (Qiagen Inc., Germantown, MD, USA). Purified DNA was submitted to Integrated Microbiome Resource (Dalhousie University, Halifax, NS) for amplicon sequencing of the bacterial 16S rRNA V6-V8 region (B969F=5’-ACGCGHNRAACCTTACC-3’ and BA1406R = 5’-ACGGGCRGTGWGTRCAA-3’) and fungal ITS2 region (ITS86(F) = 5’-GTGAATCATCGAATCTTTGAA-3’ and ITS4(R) = 5’-TCCTCCGCTTATTGATATGC-3’) on the Illumina MiSeq platform (Illumina, San Diego, CA, USA) using a 2 × 300 paired-end kit.

### 2.3. Greenhouse reciprocal transplant experiment

To disentangle the relative contribution of soil physicochemical factors and resident soil microbiota in shaping the rhizosphere microbiome, we conducted a reciprocal transplant greenhouse experiment using contrasting soils and inoculum. This complementary experiment was performed to evaluate the contribution of soil properties and soil microbial inoculum in shaping the resulting microbial community under uniform temperature and moisture conditions. A subset of ten barley lines was selected to compare those that were well adapted or un-adapted based on their relative yield performance in the 2021 swMT and cntrMT field studies (Ewing et al., 2023). Three lines each were selected based on performance at a single location (cntrMT or swMT), and four lines were selected that performed consistently across both locations.

The selected barley lines were grown under four soil conditions consisting of soil and inoculant from swMT or cntrMT. Each pot contained 80% by volume of pasteurized field soil from either swMT or cntrMT to represent base soil. Soil was pasteurized by 3 repetitions of steaming at 160F for 1.5 hours. Each repetition was conducted 24 hours apart. Ten percent by volume was non-pasteurized soil which served as the microbial inoculum. This resulted in four treatments; swMT soil with smMT inoculum, swMT soil with cntrMT inoculum, cntrMT soil with cntrMT inoculum, and cntrMT soil with swMT inoculum. The final ten percent of each pot by volume contained pasteurized sand to increase water infiltration and pore space for roots. Each line and soil combination was replicated six times and pots were watered every two days. Twice a week during watering, Peters Professional 20-20-20 fertilizer G99290 (ICL, St. Louis, MO) was added to the water at a concentration of 200 ppm nitrogen. Three replicates were removed at stem elongation to collect rhizosphere samples, and three replicates were grown to maturity and dry weight of the above ground biomass was measured. Rhizosphere extractions, DNA purification, and sequencing were performed as described above in Section 2.2. Only bacterial sequencing was performed because soil mixing and the short duration of the experiment was expected to disrupt mycorrhizal associations and confound interpretation of the fungal data (Gorzelak et al., 2015).

### 2.4. Sequencing data analysis

Quality filtering, sequence variant assignment, and alpha and beta diversity analysis were performed in R (Team, 2024). Reads were trimmed to remove the first 5 bases and truncated at 255 bp to remove low quality bases. Only forward reads were used for downstream analysis due to the low quality of reverse reads. After dereplication, sequence tables from each location-year were combined for chimera removal and taxonomy assignment using Dada2 (Callahan et al., 2016). Fungal ITS reads were filtered to remove reads < 50 bp. Bacterial ASV taxonomic classification was performed using a naive Bayes classifier pre-trained on the weighted Silva 138.1 database with a 99% identity threshold for bacterial 16S amplicons and the Unite version 8.3 fungal taxonomy database was used for ITS amplicons (Nilsson et al., 2019; Quast et al., 2013). Alpha diversity was estimated using Shannon’s diversity index after rarefying data to 10,000 reads per sample to account for uneven sequencing depth and Venn diagrams were used to visualize the number of shared and unique ASVs among locations.

### 2.5. Statistical Analysis

The effect of location, year, genotype (barley line), and interactions between these factors, on rhizosphere community composition was assessed using permutational multivariate analysis (PERMANOVA) with 9999 permutations on the Bray-Curtis dissimilarity matrix. Differences in community composition between location-years was assessed visually using constrained analysis of principal coordinates with Bray-Curtis dissimilarities using the vegan R package (Oksanen et al., 2018) with the soil parameters nitrate, potassium, organic matter, and pH as constraints.

To compare soil environments in the greenhouse we performed a constrained analysis of principal coordinates (CAP) using the weighted unifrac distances. Distances were constrained by base soil and inoculant, and the model was tested using PERMANOVA. A Random Forest model, implemented in the microeco (Liu et al., 2021a) R package, was used to identify taxa that were responsive to each base soil and inoculum combination (Breiman, 2001). To infer putative bacterial functions in the greenhouse study, predictive functional profiling of the bacterial community was performed using Functional Annotation of Prokaryotic Taxa (FAPROTAX) (Louca et al., 2016). Differences in plant biomass between treatments were evaluated using a linear mixed-effects model with base soil, inoculant, and barley genotype as random effects. Post-hoc pairwise comparisons of the estimated marginal means (EMMs) were conducted using Tukey’s Honestly Significant Difference (HSD) test. Statistical significance was evaluated at an alpha level of 0.05. To visualize group differences, a compact letter display (CLD) was generated. Statistical analyses were performed using the emmeans (Lenth, 2023) and multcomp packages (Hothorn et al., 2008).

### 2.6. Variance partitioning

Variance partitioning was used to calculate the proportion of total variance (termed “variance scores”) attributed to potential explanatory factors for rhizosphere taxa as previously described in (Meier et al., 2021; Tedersoo et al., 2014). Variance partitioning was performed at the ASV level with center-log-transformed relative abundances to estimate the contribution of environment (location-year) and soil physical and chemical parameters to changes in microbiome composition in rhizosphere. We used a 5% variance partition threshold to filter meaningful factors as previously described (Meier et al., 2021). Mixed effects models were used to assess fixed quantitative effects such as soil chemistry (NO_3_-N, OM, K, and pH) while random effects included location-year, genotype, and block nested within location-year. The final linear model was

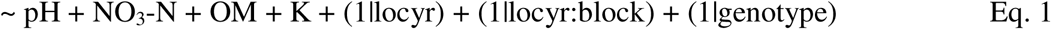

The fraction of total variance explained by each factor and total number of ASVs above the 5% threshold was then plotted to estimate the contribution of each factor to changes in microbiome composition in rhizosphere.

### 2.7. Ecotypes and community assembly

To explore the effect of environmental factors in shaping specific components of the rhizosphere communities, we defined four ecological strategies of abundant, rare, generalist, and specialist following the method of Riddley et., al. (Riddley et al., 2025). Abundant taxa were defined as ASVs with a mean relative abundance across all samples > 0.1% which was within the outlier distribution (Fig. S2) as commonly used in other studies (Liao et al., 2017; Liu et al., 2015). Rare taxa were defined as ASVs below the median of the mean relative abundance of all samples (Liao et al., 2017; Riddley et al., 2025).

Habitat generalists and specialists were defined based on Levins niche breadth which was calculated using Eq. 2 for each ASV (Levins, 1968):

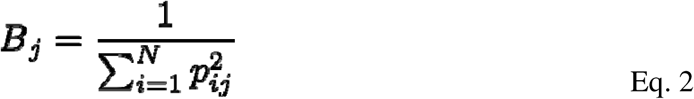

where *B_j_* is the niche breadth and *P_ij_* is the relative abundance of species *j* in a given habitat *i*. A permutation test was performed to quantify the degree to which *P_j_* deviates from expected values as previously described (Logares et al., 2013). Shannon’s diversity index was also calculated for habitat generalists and specialists.

Bacterial and fungal taxa that were over- or underrepresented in each ecotype were compared using a binomial distribution model-based enrichment index as previously described (Riddley et al., 2025). This enrichment index was used to identify over or underrepresented taxa between locations and to compare bulk soil and rhizosphere across all locations. To assess the plant genetic impact on taxa representation, enrichment was also compared among barley subpopulations. These subpopulations were determined by first calculating the genetic distance matrix for the full population then conducting a Principal Component Analysis (PCA) using the R-package “SNPRelate” followed by a Hierarchical Clustering of Principal Components (HCPC) using the R-package “FactoMineR” (Lê et al., 2008; Zheng et al., 2012) (Fig. S1).

Processes governing community assembly were inferred using the beta nearest taxon index (βNTI) and Raup-Crick based Bray-Curtis dissimilarity (RC_Bray_) null-model metrics as described in other studies (Ohigashi et al., 2025; Peng et al., 2024). βNTI calculations were performed with iCAMP as implemented in the microeco R package (Liu et al., 2021a; Ning et al., 2020). The βNTI metric compares observed phylogenetic turnover between communities with turnover expected under a null model where values of βNTI > 2 were classified as heterogeneous selection, indicating phylogenetic turnover was significantly greater than expected and βNTI < -2 represented homogeneous selection where phylogenetic turnover was significantly less than expected under the null models (Peng et al., 2024). Dispersal limitation was quantified as the fraction of pairwise comparisons with |βNTI| < 2 and RC_Bray_ > 0.95 while homogenizing dispersal was defined as the fraction of pairwise comparisons with |βNTI| < 2 and RC_Bray_ < -0.95 (Ohigashi et al., 2025). Pair-wise comparisons with |βNTI| < 2 and |RC_Bray_| < 0.95 were defined as weak selection or dispersal, or drift (Ohigashi et al., 2025). Inferred assembly processes were compared among location-years, between bulk soil and rhizosphere communities, and among barley genetic clusters.

## 3. Results

### 3.1. Diversity and composition of microbial communities

We characterized the rhizosphere microbial communities of the S2MET barley population and bulk soil using 16S and ITS amplicon sequencing across seven location-years. The sites included two locations in Montana and one in South Dakota which represent barley growing regions in the northern Great Plains. A contrasting environment was also compared near Oahu, Hawai’i, which lies outside of the adapted range where the S2MET barley population was developed. Following quality filtering and chimera removal, a total of 2,086 16S samples with an average read depth of 34,037 reads per sample and 1,432 ITS samples with an average read depth of 20,060 reads per sample were retained. The distribution of mean relative abundance of ASVs was consistent across bacterial and fungal communities (Fig. S2).

Bacterial and fungal alpha diversity, as measured by Shannon’s diversity index, were not significantly different (p > 0.05) across locations or years (Fig. 1 A and 1 B). Compositionally, bacterial communities were predominantly composed of *Actinobacteriota* and *Proteobacteria* at the phylum level. Principal component analysis showed that bacterial and fungal communities in HI differed from those in the adapted range of barley (Fig. S3). In locations within the adapted range of barley, *Actinobacteriota* tended to dominate, while in Hawai’i, *Proteobacteria* was most abundant (Fig. 1 C). *Ascomycota* was the predominant fungal phyla across all location-years (Fig. 1 D). Results of the PERMANOVA show significant (p < 0.001) main effects for location and year. Barley genotype (line) was significant (p < 0.001) for fungi but not for bacteria. A significant (p < 0.001) location x year interaction was also observed (Table S2), thus subsequent beta diversity analyses were based on location year.

**Fig. 1.**
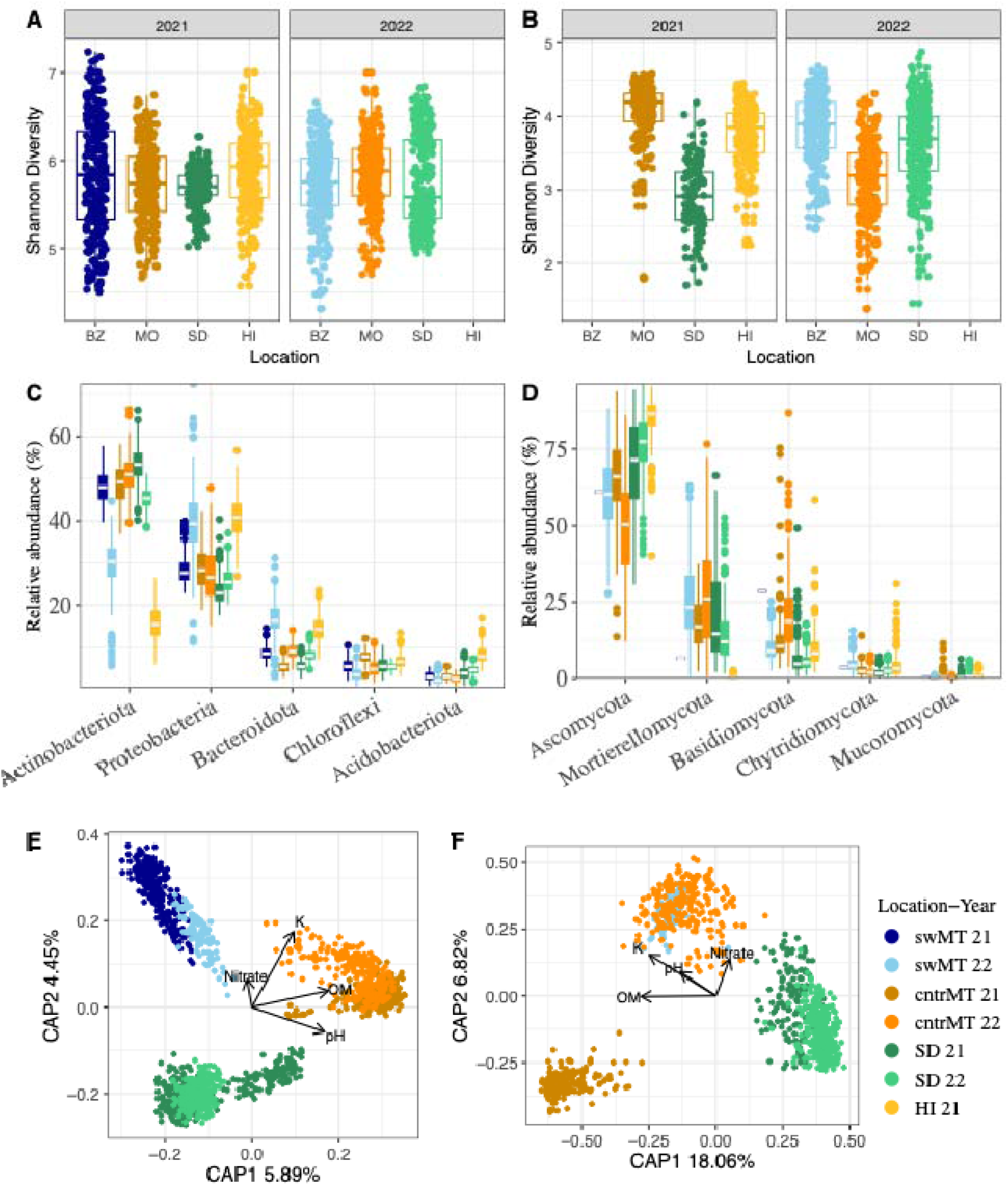
Summary of rhizosphere bacterial and fungal diversity and composition across location-years. Shannon’s diversity for bacterial (A) and fungal (B) communities. Relative abundance of major bacterial (C) and fungal (D) phyla across location-years. Constrained ordination of bacterial (E) and fungal (F) communities within barley growing regions with vectors for soil factors. Variation between location-years was significant and greater than variation within location-year for bacterial (PERMANOVA, p = 0.0002, F = 34.17) and fungal (PERMANOVA, p = 0.0002, F = 121.02) communities. Study was only performed one year (2021) in Hawaii and was not included in the constrained ordination (See Fig. S3). Missing fungal ITS data in 2021 at the swMT location was due to most reads failing to pass the quality filtering threshold.

Results of the constrained principal coordinates analysis showed that nitrate, potassium, pH, and organic matter concentrations explained most of the variation between location years and were strongly associated with the cntrMT location (Fig. 1 E). However, the first two principal coordinates only explained 7.31% of the variability in bacterial communities, indicating high variation within site-year. In contrast, fungal community diversity showed relatively greater variability across location-years with 23.1% of the variability explained by the first two principal coordinates (Fig. 1 F).

### 3.2. Variance partitioning reveals soil and location-year effects

Variance partitioning revealed location-year as the factor that explained the most variance. Within bacterial communities, a total of 157 ASVs were associated with location-year (Fig. 2A) while 115 ASVs were responsive to soil factors, with 59, 40, and 37 ASVs responsive to pH, organic matter, and potassium, respectively. Fungal variance was also largely associated with location-year, with 122 ASVs above the 5% variance threshold responsive to location-year and 78% of variance explained by this factor (Fig. 2B). A similar response was observed in fungi where ASVs were responsive to soil factors, with 39, 28, and 22 ASVs responsive to pH, organic matter, and potassium, respectively. In addition, 59 bacterial and 43 fungal ASVs were associated with a block x location-year interaction.

**Fig. 2.**
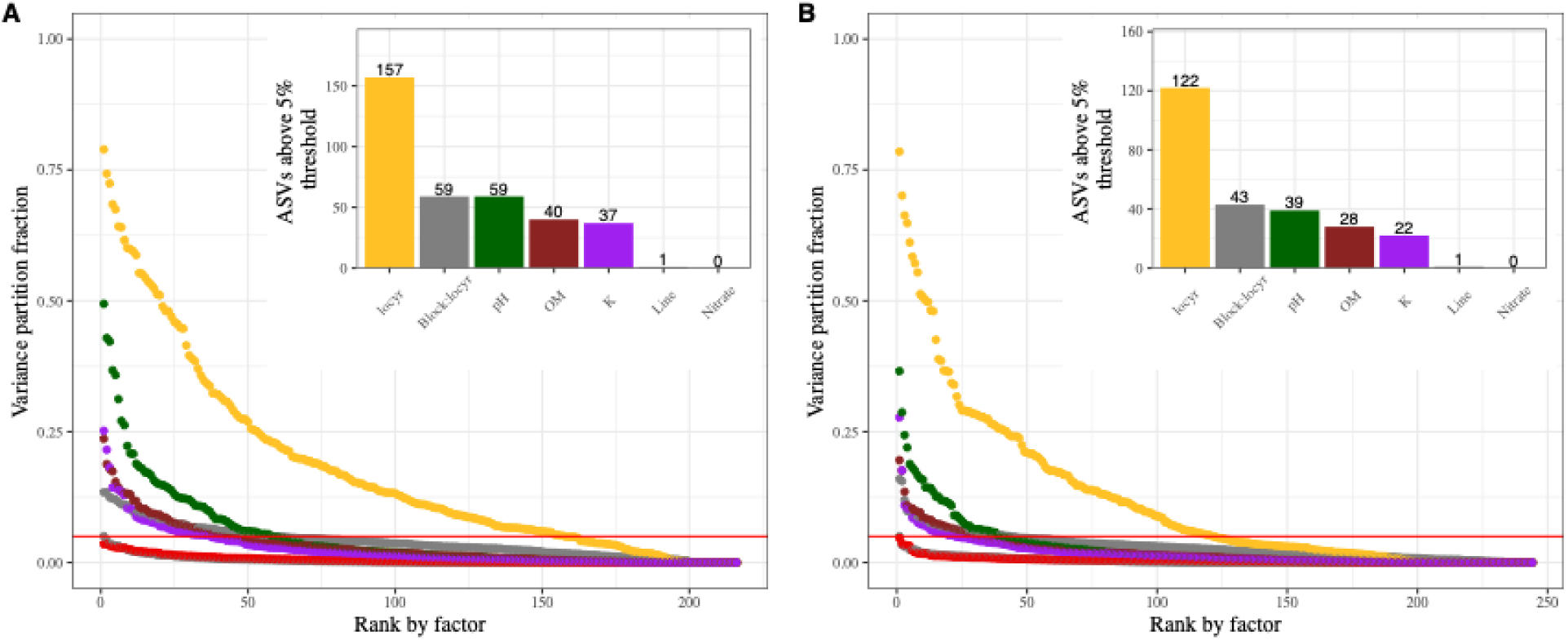
Variance partitioning for (A) bacterial and (B) fungal genus abundances. A linear model was calculated per genus as fixed qualitative and random categorical variables (∼ pH + OM + K + Nitrate + (1|locyr) + (1|Block)).

### 3.3. Ecotypes and community assembly

Rhizosphere bacterial and fungal ASVs were classified, based on ecological traits, as abundant, rare, generalists, or specialists. Among abundant bacterial phyla, *Actinobacteriota* was significantly (p < 0.05) enriched, while *Bacteriodota* and *Chloroflexi* were depleted (Fig 3 A). In contrast, no rare ecotypes were enriched or depleted. *Actinobacteriota* was enriched among generalists while *Verrucomicrobiota* was underrepresented. Among specialists, *Actinobacteriota* and *Proteobacteria* ASVs were enriched while *Verrucomicrobiota* was depleted (Fig. 3 A).

**Fig. 3.**
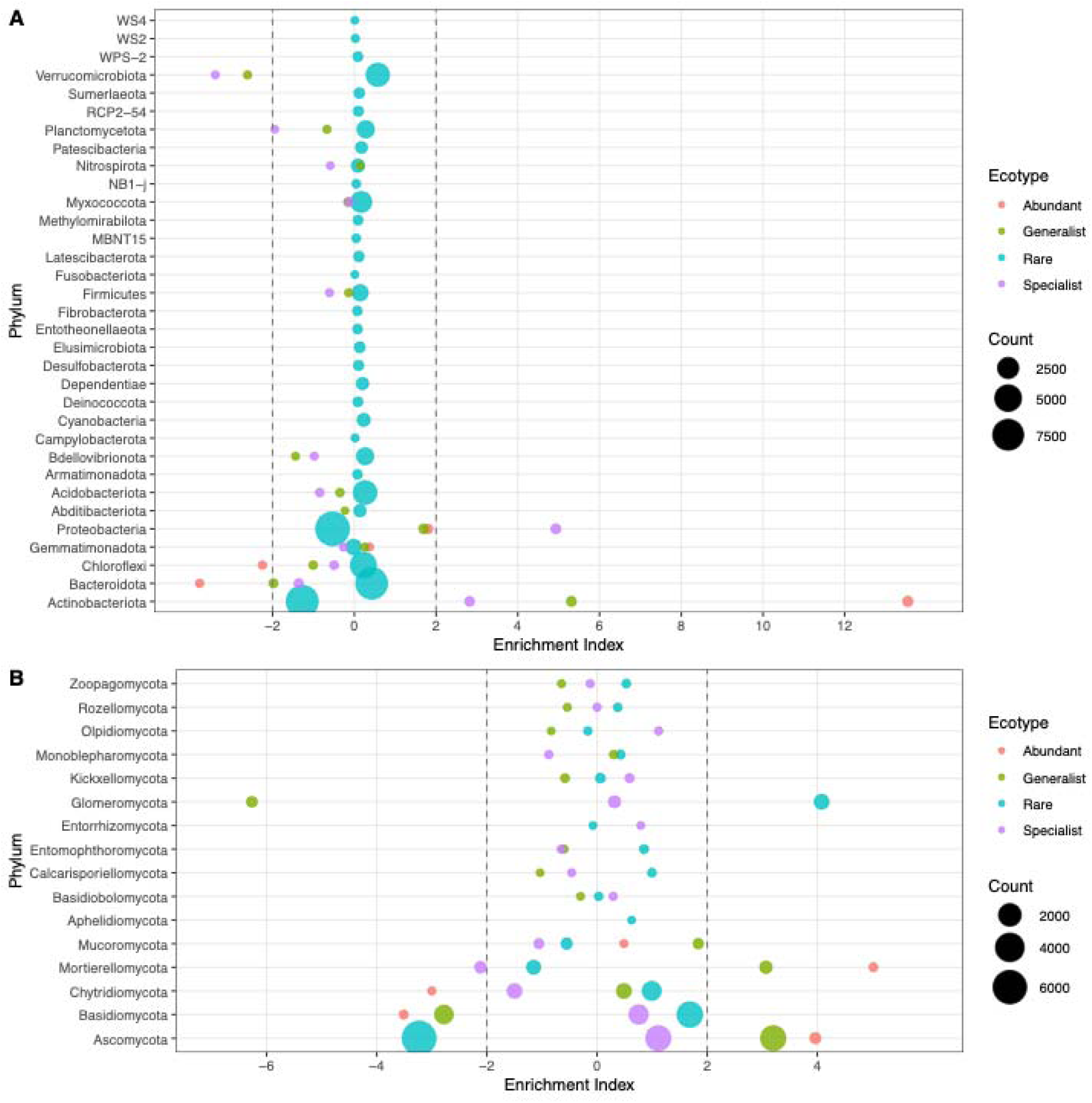
Enrichment analysis at the phylum level comparing ecotype for rhizosphere bacterial (A) and fungal (B) communities. Circle size indicates the number of ASVs representing a phylum for each cluster. Phyla with an enrichment index >2 or < -2 indicate significant over or underrepresentation within each cluster.

Among abundant fungal taxa, Ascomycota and Mortierrellomycota were enriched, while Basidiomycota and Chytridiomycota were depleted (Fig. 3 B). In contrast to bacteria, several rare fungal phyla had significant (p < 0.05) enrichment index values. Glomeromycota was enriched, while many ASVs associated with the phylum Ascomycota were underrepresented. Among generalist taxa, Ascomycota and Mortierellomycota were overrepresented and Basidiomycota and Glomeromycota underrepresented. Mortierellomycota was significantly underrepresented among fungal specialists (Fig. 3 B).

Enrichment analysis was performed to compare rhizosphere communities to bulk soil regardless of location. Bacteriodota, Bdellovibrionota, Cyanobacteria, and Verrucomicrobiota were significantly (p < 0.05) enriched in the rhizosphere while Actinobacteriota and Acidobacteriota were depleted relative to bulk soil (Fig. S4 A). In contrast, no fungal phyla were significantly enriched in the rhizosphere relative to bulk soil (Fig. S4 B). Enrichment analysis also identified multiple phyla that were over- or underrepresented based on location. HI had the greatest number of over- and underrepresented bacterial taxa. Notably, four phyla had enrichment indices < -6 which was much lower than index values for locations within barley growing regions. (Fig. 4 A). Actinobacteriota and Proteobacteria were highly overrepresented in HI compared to locations within barley growing regions. Fungal phyla were more location specific with 64% of all fungal taxa being over- or underrepresented in one or more locations (Fig. 4 C).

**Fig. 4.**
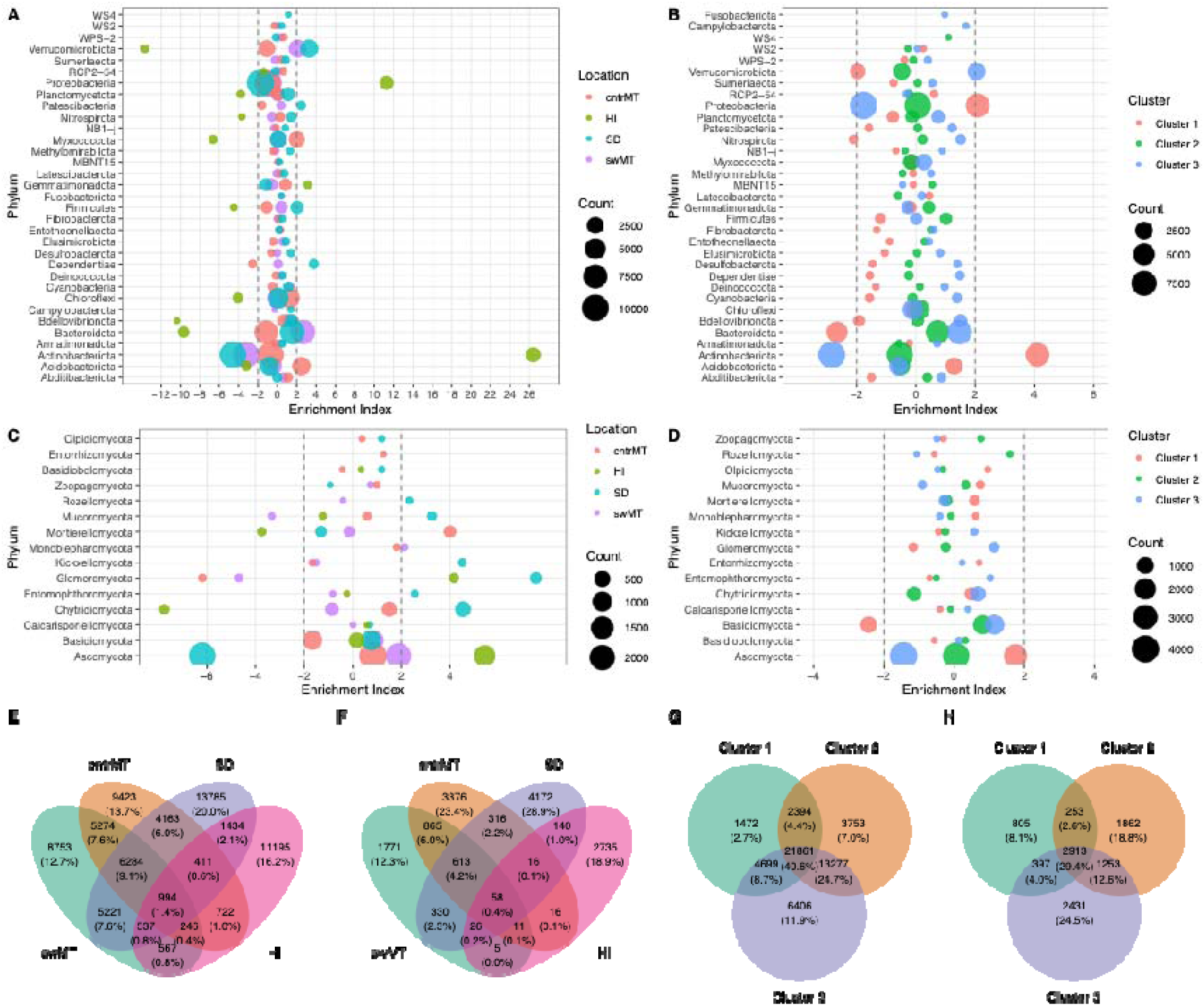
Enrichment analysis of phyla among location for rhizosphere bacteria (A) and fungi (C). Enrichment analysis at the phylum level comparing barley genotypes grouped by PCA hierarchical clustering for rhizosphere bacterial (A) and fungal (B) communities. Circle size indicates the number of ASVs representing a phylum for each cluster. Phyla with an enrichment index >2 or < -2 indicate significant over or underrepresentation within each cluster. Venn diagrams showing shared and unique rhizosphere ASVs among locations for bacteria (E) and fungi (F) and among barley genotypes grouped by PCA hierarchical clustering for bacteria (G) and fungi (H).

We next examined whether enrichment patterns differed among barley subpopulations defined by PCA-based hierarchical clustering of genotype data. In bacterial communities, Actinobacteriota and Proteobacteria were significantly (p < 0.05) enriched in cluster 1 and depleted in cluster 3 while Bacteriodota and Nitrospirota were depleted in cluster 1 (Fig. 4 B). Fungal differences among barley clusters were more limited with Basidiomycota being the only phylum depleted in cluster 1 (Fig. 4 D).

We also compared the number of unique and shared ASVs among locations and barley subpopulations. Only 1.4% of bacterial ASVs were shared among all locations while 9.1% of ASV were shared among the adapted barley regions of swMT, cntrMT, and SD, demonstrating an environmental limitation on the ability of plants to recruit rhizosphere community members outside of their typical growing region (Fig. 4 E). Fungal communities were even more location specific with only 0.4% of fungal ASVs shared among all locations, and 12.3% – 28.9% were unique to one location (Fig. 4 F). Although enrichment analyses showed relatively few phylum-level differences among barley subpopulations, ASV level comparisons showed stronger differentiation with 21.6% bacterial and 51.4% fungal ASVs that were unique to only a single barley subpopulation (Fig. 4 G, H). More ASVs were shared between cluster 2 and 3 (24.7% and 12.6% for bacteria and fungi, respectively) compared to cluster 1.

Finally, we compared inferred community assembly processes among locations, between rhizosphere and bulk soil, and among barley subpopulations. In both bacterial and fungal communities, stochastic processes, including dispersal limitation, homogenizing dispersal, and drift, accounted for around 95% of assembly processes across all location-years (Fig. S5) indicating neutral processes were the primary drivers shaping community assembly. Bacterial communities showed similar patterns when compared between bulk and rhizosphere soil and among barley gene clusters (Fig. 5). In contrast, fungal communities showed a stronger contribution of deterministic processes. Specifically, homogenizing selection, where phylogenetic turnover was lower than expected under the null model, had a greater role in the rhizosphere than in bulk soil and differed among barley subpopulations (Fig. 5A, B). Heterogeneous selection, where phylogenetic turnover was higher than expected, was the main deterministic process in clusters 1 and 2 while homogeneous selection was the dominant deterministic process in cluster 3.

**Fig. 5.**
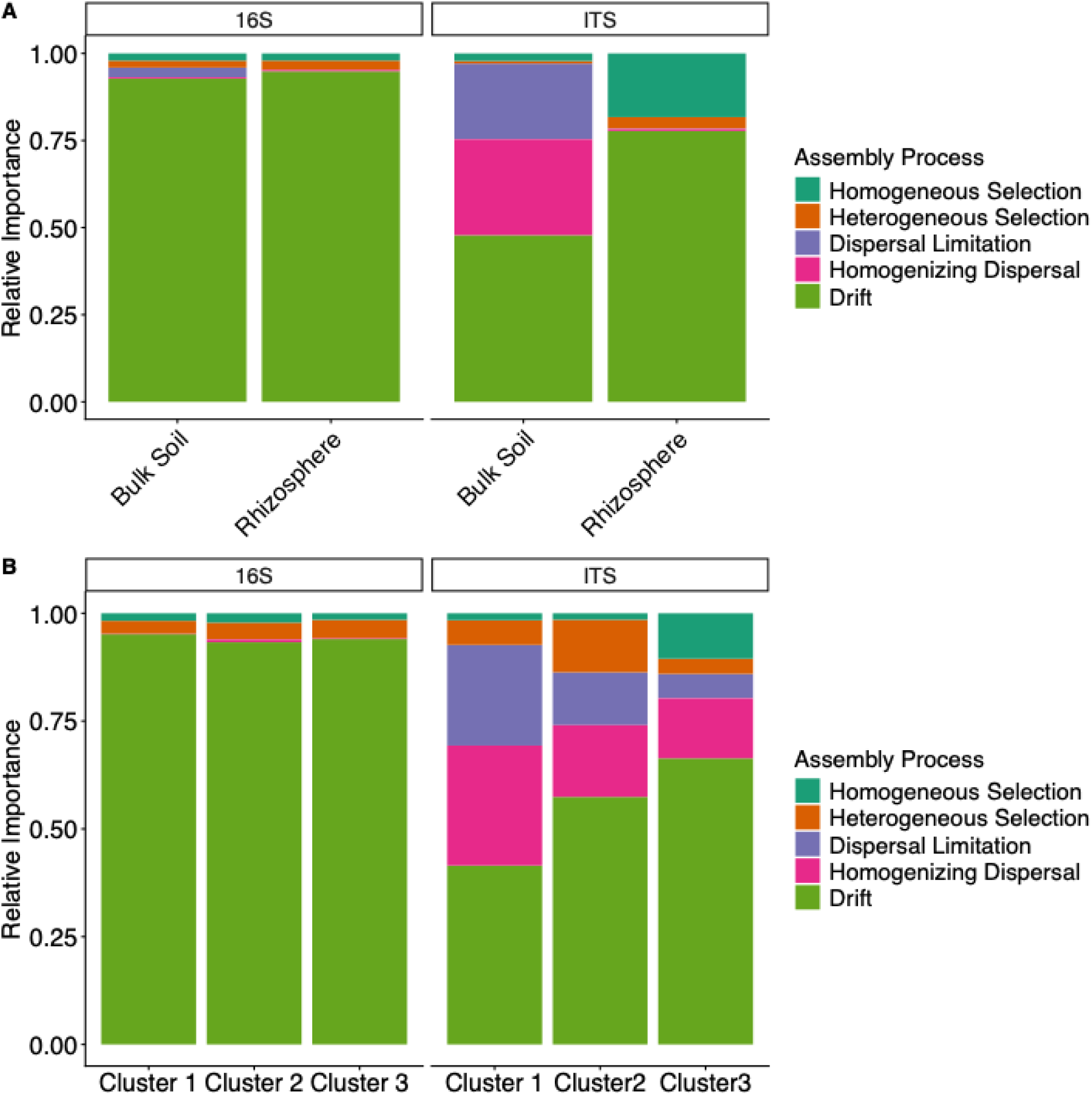
Relative importance of stochastic and deterministic ecological processes in shaping rhizosphere bacterial and fungal community assembly comparing bulk soil to rhizosphere communities (A) and hierarchical PCA clustering of barley gene loci (B). Deterministic processes include homogeneous and heterogeneous selection while stochastic processes include dispersal limitation, homogenizing dispersal, and drift.

### 3.4. Reciprocal transplant experiment

The pronounced location effects in field trials raised the question of whether these differences stem from inherent soil physicochemical properties or the resident microbial communities unique to each site. To partition the effects of the physicochemical characteristics of soils from microbial communities and evaluate the effects of these factors on plant growth, we performed a greenhouse reciprocal transplant experiment with a subset of ten S2MET barley genotypes using soils from locations with contrasting yield potentials and microbial community composition. We prepared four combinations of base pasteurized soils and inoculum from cntrMT and swMT (Fig. 6 A). Results demonstrated the distinct influence of the base soil physicochemical properties and soil inoculum in shaping rhizosphere communities. Results of the PERMANOVA showed a significant base soil, inoculant, barley genotype, and base soil x inoculant interaction (Table S3).

**Fig. 6.**
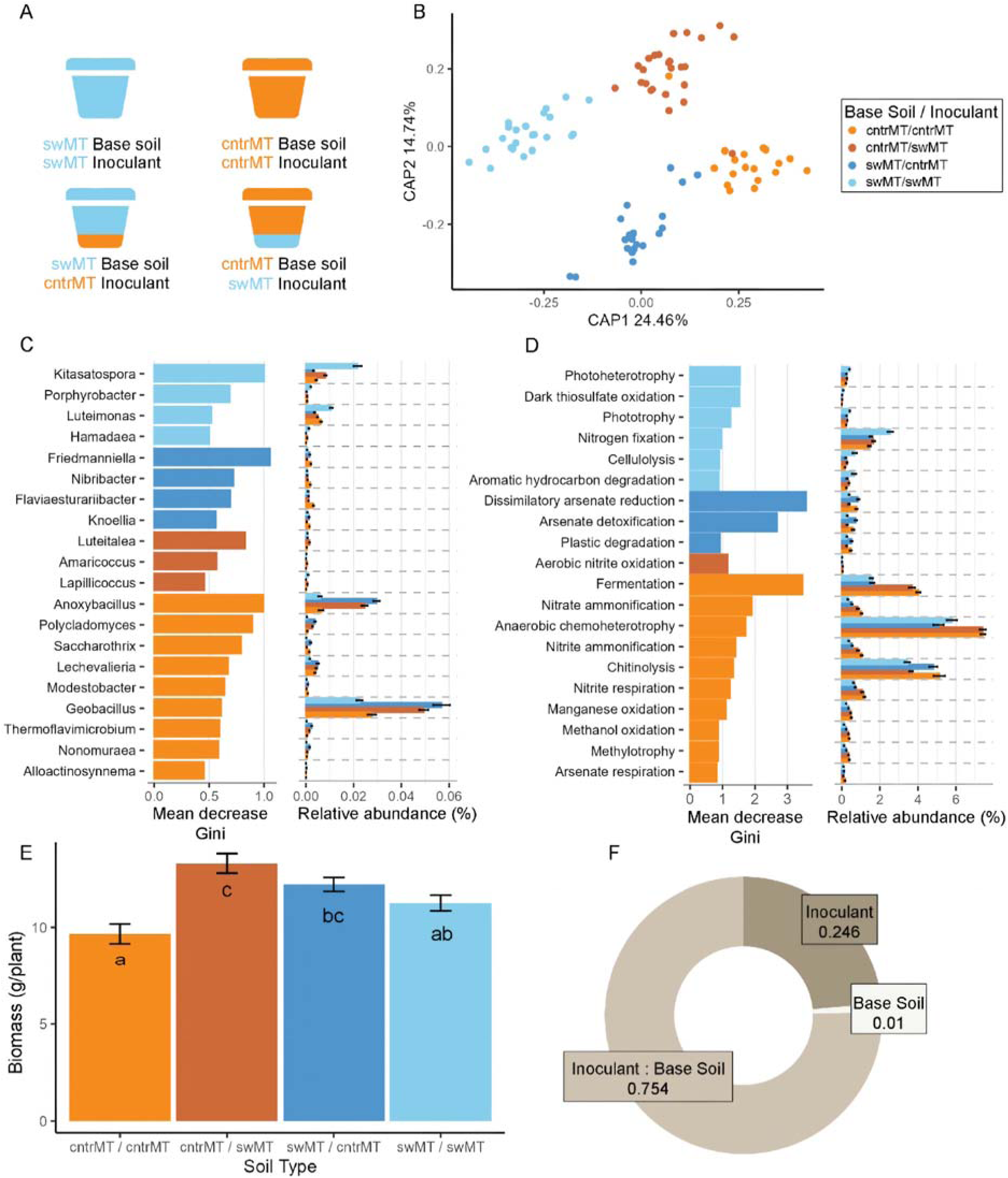
Greenhouse study evaluating the contribution of soil and inoculum on rhizosphere microbiome composition and barley biomass. (A) Schematic showing the combinations of base soil and inoculum used in the experiment. (B) Constrained analysis of principal coordinates (CAP) of the weighted unifrac distances of the rhizosphere bacterial communities based on the base soil and inoculant. Differentially abundant genera (C) and functions (D) predicted by a random forest model with a Mean Decrease Gini (MDA) threshold of 10. Functions were predicted from 16S sequences using the functional annotation of prokaryotic taxa (FAPROTAX) database. Only significant (adjusted p < 0.001) taxa are shown. (E) Barley dry biomass at harvest across soil treatments. Letters indicate significant differences at p<0.05. (F) Variance partitioning of the effects of base soil, inoculant, and their interaction on barley dry biomass. cntrMT: central Montana, swMT: southwest Montana.

Rhizosphere composition differed in response to both base soil and inoculum source (Fig. 6 B). The genus *Kitasatospora* was enriched with swMT inoculum, regardless of the base soil (Fig. 6 C). In contrast, the genera *Anoxybacillus* and *Geobacillus* were most abundant when soils from one location were paired with inoculum from the other location while *Luteimonas* was more abundant when soils were inoculated with microorganisms from the same location.

In contrast to the response of specific taxa, differences in the abundance of predicted functions were driven primarily by the base soil source. Predicted functions associated with nitrogen fixation, cellulolysis, and aromatic hydrocarbon degradation were more abundant in swMT/swMT relative to the other treatments (Fig. 6 D). Chitinolysis was more abundant with cntrMT inoculum compared to swMT while functions related to fermentation and anaerobic chemoheterotrophy were more abundant in cntrMT soil, regardless of inoculum source. Manganese oxidation was more abundant in cntrMT soil (Fig. 6 D). Soil tests showed that manganese was > 2.5-fold higher in cntrMT than swMT (Table S1), revealing a potential base soil-inoculum interaction resulting in greater abundance of manganese oxidation functions with cntrMT soil regardless of inoculum source.

We examined whether these bacterial community-shaping contexts of base soil and inoculum interact to drive differences in plant growth. Plant biomass varied due to base soil and inoculum source (Fig. 6 E-F). Interestingly, the combination of cntrMT base soil with swMT inoculant resulted in the greatest biomass (Fig. 6 E), while base soils with inoculum from the same location had the lowest biomass. Variance partitioning of the effects of base soil, inoculant, and their interaction on barley dry biomass indicated that base soil only explained 1% of the variance while inoculant and inoculant x base soil interaction explained 24.9% and 75.4%, respectively (Fig. 6 F).

## Discussion

Rhizosphere microorganisms play a fundamental role in plant nutrient acquisition, abiotic stress protection, and pathogen suppression. Understanding the drivers of recruitment and the role of abiotic and biotic factors in shaping community assembly is important for future efforts to enhance crop productivity and sustainability by identifying new targets for crop breeding and improving sustainable management strategies. By leveraging a multi-environment field study and a controlled greenhouse reciprocal transplant experiment, we disentangled the contributions of location, soil properties, the resident microbiota, and barley genotype to rhizosphere assembly.

Environment exerted a greater influence on rhizosphere communities than host genotype which supports hypothesis (i). Across barley growing regions of Montana and South Dakota, bacterial communities were broadly consistent and were predominantly Actinobacteriota and Proteobacteria which is consistent with other studies (Maestre et al., 2015; Riddley et al., 2025) while fungal communities were dominated by the phylum Ascomycota, consistent with global soil surveys (Egidi et al., 2019). Importantly, by including a contrasting environment outside the adapted range of barley, we demonstrated an extreme case of environmental filtering and a limit to plant recruitment of soil microorganisms, supporting hypothesis (ii) that a novel environment yields a divergent rhizosphere community.

Variance partitioning analysis further quantified this hierarchy of drivers. Location-year consistently explained the largest share of variation in microbial community composition across taxa, whereas specific soil physicochemical factors (e.g. pH, nutrients) had smaller, though measurable, contribution. The strong location-year effect may have been due in part to drought conditions in the Montana locations. In swMT in 2021, May – August precipitation was 17% below the 100-year average while in cntrMT, May – August precipitation was 24% and 50% below the 100-year average in 2021 and 2022, respectively Table S1). Community assembly metrics reinforced the predominance of broad-scale stochastic processes. The phylogenetic turnover analysis (βNTI and RC metrics) indicated that over 90% of community differences across samples were attributable to neutral or stochastic assembly (random drift and dispersal), with only a small proportion of instances showing evidence of deterministic selection. This is consistent with other studies which have shown stochastic processes dominate (Chen et al., 2026; Ma et al., 2025). For fungal communities, there was a slightly higher deterministic signal than for bacteria, yet in both cases stochastic processes were dominant. These results suggest that broad-scale climate and edaphic conditions that vary annually play a greater role in shaping community structure than deterministic soil chemistry filtering. Soil microbial communities are sensitive to changes in temperature and moisture, and these factors also alter host plant physiology (Meier et al., 2021) which may have an indirect effect on microbial recruitment. One limitation of these findings was that individual climate factors could not be included in variance partition models because conditions such as precipitation and temperature were representative of each location-year and were not measured at a barley line or block level.

Despite the predominant influence of environmental conditions and stochastic assembly processes, we observed evidence of host genotype filtering on specific taxa, consistent with our hypothesis (i). Multiple ASVs belonging to the phyla Actinobacteriota and Proteobacteria were significantly (p < 0.05) enriched in genotype cluster 1 and depleted in cluster 3 while Bacteriodota and Nitrospirota were depleted (Fig. 4 B). Enrichment analysis identified only one fungal phylum, Basidiomycota, which was depleted in cluster 1 (Fig. 4 D). However, these genotype-associated patterns were not uniform across all environments, nor did they manifest as broad shifts in whole-community composition, as evidenced by the negligible genotype effect in overall β-diversity analyses. The significance of these genotype-specific ASVs must be interpreted cautiously since they may reflect low-abundance taxa that are strongly influenced by chance colonization events. Overall, this finding provides additional support for stochastic processes being the primary driver of community assembly. Within this context, barley genetics appear to fine-tune the microbiome by altering the prevalence of specific taxa rather than changing the overall community structure. Our findings align with a growing body of evidence that plant host effects are context-dependent and genotype-driven microbial associations often vary by environment and may be reproducible only under certain conditions.

A strength of this research is the combination of field trials with a greenhouse reciprocal transplant experiment, which allowed for a more mechanistic understanding of how abiotic and biotic soil components influence rhizosphere assembly. While soil microorganisms are often considered part of the environment (Blouin et al., 2026), our study provides experimental evidence that soil physicochemical factors and resident microbial community independently shape the rhizosphere community composition. Constrained ordination confirmed that differences in base soil explained a large portion of variation in rhizosphere microbiomes, while the source of the microbial inoculum also had significant though smaller effect. Differences in individual taxa were driven primarily by source inoculum, supporting the idea that rhizosphere communities are shaped in part by the available soil inoculum. In contrast to taxonomy, differences predicted functional abundances were driven primarily by base soil which is consistent with the expectation of functional redundancy among taxa and reflects a functional response to soil chemistry such as the abundance of C, N, and other nutrients (Louca et al., 2018). Barley genotype effect was also significant, but the proportion of variance explained was small suggesting a limited impact relative to soil and inoculum (Table S3). These results support the field observations that differences in soil chemistry and climate across locations determine which microbes can thrive, and this pool of microbes present in the soil further constrains which taxa are recruited by roots. The greenhouse experiment reinforces our interpretation that environment-driven effects on microbiome composition arise both from abiotic filtering and biotic filtering. Coupled with field data, this provides causal evidence that the barley microbiome is shaped by an interaction between soil factors and microbial inoculum, with host genetics modulating recruitment within that framework. One limitation of this study is that functional predictions were constrained by the genus level or higher taxonomic resolution of the sequencing data. Microbial function can vary even at the species or strain level and predictive functional profiling cannot resolve strain level variations or identify functions if the pathway annotations are poor (Langille et al., 2013). While this approach is useful for hypothesis generation, metagenomic or metatranscriptomic studies are needed to validate predicted functions.

## Conclusions

By explicitly assessing environment, soil, and plant genetic factors together, our study presents a hierarchical model of rhizosphere assembly in barley. Broad-scale environmental factors emerge as the top-tier drivers, defining the baseline community composition and diversity. Within this environmental context, the rhizosphere community is further shaped by local soil chemical properties and the existing soil microbial community. Finally, plant genotype introduces a fine-scale filter by altering the relative abundance of select microbial taxa without drastically changing the overall community structure. These insights confirm our initial hypotheses and refine our understanding of how adaptation to local conditions is reflected in the crop microbiome. Barley largely recruits a broadly similar microbiota across adapted environments, whereas in a climatically extreme, poorly adapted site, environmental filtering dominates to yield a novel community. Meanwhile, plant genetic diversity in barley still has consequential effects on specific microbes, which may have functional relevance.

The predominantly stochastic nature of community assembly we observed demonstrates that microbiome outcomes in field conditions can be variable, posing challenges for harnessing these interactions reliably. However, the detection of genotype-associated microbial differences within each environment suggests that targeted breeding or management to favor certain beneficial taxa must be pursued with recognition of strong environmental context. The primary constraint to achieving these goals is a lack of a mechanistic understanding of specific plant-microbe interactions and future research is needed to build on these findings by integrating barley genomics and functional microbial traits to identify mechanisms underlying these interactions.

## Acknowledgements

This work was supported by the United States Department of Agriculture (USDA) National Institute of Food and Agriculture (NIFA) Agriculture and Food Research Initiative (AFRI) award number 2020-67014-32138. Computational efforts were performed on the Tempest High Performance Computing System, operated and supported by University Information Technology Research Cyberinfrastructure at Montana State University. The authors thank the Hawaii Agricultural Research and the Montana State Agricultural Experiment Stations for support. Mention of trade names or commercial products in this publication is solely for the purpose of providing specific information and does not imply recommendation or endorsement by the U.S. Department of Agriculture. The USDA is an equal opportunity provider and employer.

## CRediT authorship contribution statement

**Erik Z. Killian**: Investigation, Data curation, Formal analysis, Visualization, Writing – original draft, Writing – reviewing & editing. **Jessica L. Williams**: Investigation, Writing – reviewing & editing. **Anna Halpin-McCormick**: Formal analysis, Writing – reviewing & editing. **Patrick M. Ewing**: Conceptualization, Funding acquisition, Resources, Methodology, Supervision, Investigation, Data curation, Writing – reviewing & editing. **Michael B. Kantar**: Conceptualization, Funding acquisition, Resources, Methodology, Supervision, Investigation, Data curation, Writing – reviewing & editing. **Jennifer Lachowiec**: Conceptualization, Funding acquisition, Resources, Methodology, Supervision, Investigation, Writing – reviewing & editing. **Jamie D. Sherman**: Conceptualization, Funding acquisition, Methodology, Resources, Investigation, Writing – reviewing & editing. and **Jed O. Eberly**: Conceptualization, Funding acquisition, Project administration, Methodology, Resources, Supervision, Investigation, Data curation, Validation, Writing – reviewing & editing.

## Declaration of generative AI and AI-assisted technologies in the manuscript preparation process

Statement: During the preparation of this work, Microsoft CoPilot 365 was used to assist with language refinement, structural organization, and clarity of expression. After using this tool/service, the author(s) reviewed and edited the content as needed and take(s) full responsibility for the content of the published article.

## Data Availability

The raw sequence data are deposited in the NCBI Sequence Read Archive (https://www.ncbi.nlm.nih.gov/bioproject/PRJNA909304). All read processing steps and custom scripts are available on GitHub (https://github.com/erikthekillian/). Processed data, R code, and metadata supporting this manuscript are available from Dryad: DOI: 10.5061/dryad.jh9w0vtsf.

