## Supplemental Material for "Environment and plant genetics shape barley rhizosphere microbiome structure across contrasting locations"

Erik Z. Killian^1^, Jessica L. Williams^1^, Anna Halpin-McCormick^2^, Patrick M. Ewing^3^, Michael B. Kantar^2^, Jennifer Lachowiec^1^, Jamie D. Sherman^1^, and Jed O. Eberly^4*^

^1^Plant Sciences & Plant Pathology Department, Montana State University, Bozeman, Montana, USA

^2^Department of Tropical Plant and Soil Science, University of Hawaii at Manoa, Honolulu, Hawaii, USA

^3^USDA-ARS Food Systems Research Unit, Burlington, Vermont, 05405, USA

^4^Central Agricultural Research Center, Montana State University, Moccasin, Montana, USA

**Table S1:** Climate and soil properties at each study location.

|  | Location | | | |
| --- | --- | --- | --- | --- |
|  | Bozeman, MT | Moccasin, MT | Brookings, SD | Kunia, HI |
| Coordinates | 45.673484, -111.155487 | 47.06242, -109.94765 | 44.350227, -96.806430 | 21.384253, -158.038122 |
| Mean annual rainfall (mm) | 411.2 | 388.6 | 640 | 813.1 |
| Mean growing season temperature (°C) | 13.8 | 13.3 | 15.4 | 22 |
| Frost free days | 110 | 112 | 145 | 365 |
| Cumulative precipitation May 1-August 31 2021 | 172 | 171 | 204 | 296.7 |
| Cumulative precipitation May 1-August 31 2022 | 198 | 115 | 360 | NA |
| Soil classification | Amsterdam silt loam | Danvers-Judith clay loam | Barnes Lay Loam | Molokai silty clay loam |
| Taxonomic Class | Fine-silty, mixed, superactive, frigid Typic Haplustolls | Fine-loamy, carbonatic, frigid Typic Calciustolls | Fine-loamy, mixed, superactive, frigid Calcic Hapludolls | Very-fine, kaolinitic, isohyperthermic Typic Eutrotorrox |
| Soil depth (cm) | 40-121 | 30-60 | >200 | 38-127 |
| pH (in water) | 6.1 | 7.9 | 6.7 | 6.8 |
| OM (%) | 3.45 | 4.6 | 3.5 | 2.6 |
| Nitrate-N (ppm) | 6.2 | 7 | 4.7 | 2.1 |
| P_2_O_5_ (ppm, Mehlich) | 50.7 | 35.1 | 8.4 | 6 |
| K_2_O (ppm) | 277.9 | 826.8 | 127 | 380 |
| Sulfate-S (ppm) | 5.6 | 12.6 | 7 | 21 |
| Ca (ppm) | 2111.5 | 4469 | 2463 | 1066 |
| Mg (ppm) | 460.6 | 193.2 | 377 | 329 |
| Mn (ppm) | 19.8 | 48.8 | 13 | 149 |

**Table S2:** Pairwise permutational analysis of variance (PERMANOVA) for multivariate (β-diversity) based on the Bray-Curtis dissimilarity matrix. Group significance is shown for location, year, and barley line and interactions between these factors.

| 16S | Df | SumOfSqs | R2 | F | Pr(>F) | |
| --- | --- | --- | --- | --- | --- | --- |
| Location | 3 | 50.44 | 0.07 | 45.8 | 0.001 | *** |
| Year | 1 | 3.21 | 0 | 8.75 | 0.001 | *** |
| Line | 238 | 87.05 | 0.12 | 1 | 0.606 |  |
| Location:Year | 2 | 6.79 | 0.01 | 9.24 | 0.001 |  |
| Location:Line | 697 | 234.42 | 0.31 | 0.92 | 1 |  |
| Year:Line | 235 | 93.55 | 0.13 | 1.08 | 0.001 | *** |
| Location:Year:Line | 406 | 161.35 | 0.22 | 1.08 | 0.001 | *** |
| Residual | 296 | 108.67 | 0.15 |  |  |  |
| Total | 1878 | 745.47 | 1 |  |  |  |
| ITS |  |  |  |  |  |  |
| Location | 3 | 192.62 | 0.4 | 402.36 | 0.001 | *** |
| Year | 1 | 27.09 | 0.06 | 169.79 | 0.001 | *** |
| Line | 238 | 42.47 | 0.09 | 1.12 | 0.001 | *** |
| Location:Year | 2 | 27.56 | 0.06 | 86.37 | 0.001 | *** |
| Location:Line | 615 | 104.21 | 0.22 | 1.06 | 0.008 | ** |
| Year:Line | 191 | 36.19 | 0.08 | 1.19 | 0.001 | *** |
| Location:Year:Line | 86 | 14.44 | 0.03 | 1.05 | 0.128 |  |
| Residual | 201 | 32.08 | 0.07 |  |  |  |
| Total | 1337 | 476.66 | 1 |  |  |  |

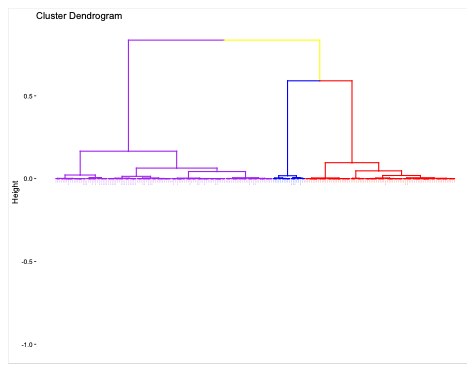

**Fig. S1**. Hierarchical clustering on principal components (HCPC) of the S2MET barley (Hordeum vulgare) mapping population designed for multi-environment genomic selection. The population consists of a training set of 183 individuals all genotyped via genotyping-by-sequencing (GBS). Principal components were computed from the genomic marker matrix using SNPRelate in R, and HCPC analysis and dendrogram visualization were conducted using FactoMineR and factoextra, respectively. Cluster height reflects dissimilarity in PC space, with shorter branch lengths indicating greater genomic similarity. Lines originate from the USDA-ARS Aberdeen, Idaho (AB), Montana State University (MT), Washington State University (WA), and North Dakota State University (N2) breeding programs, alongside advanced 2MS14 entries. Clustering patterns reveal broad separation between the 2MS14 lines and earlier-cycle program entries, with partial grouping by program of origin suggesting regional population structure within the panel.

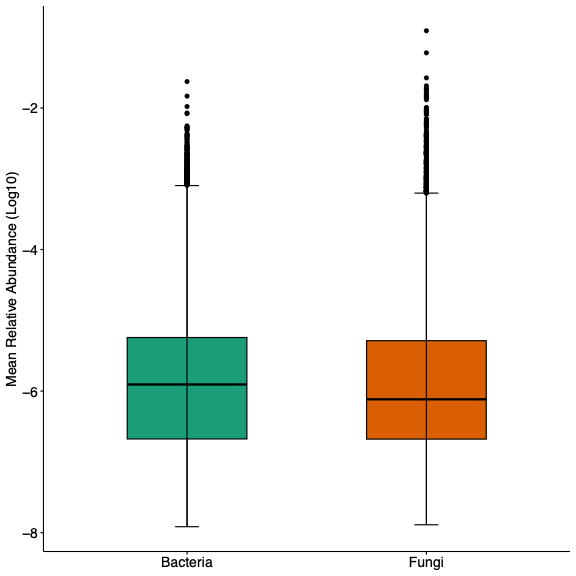

**Fig. S2.** Distribution of mean relative abundance (log10) of ASVs across bacterial and fungal samples. Box plots show the interquartile range (IQR), with the line representing the median and whiskers extending to 1.5 times the IQR.

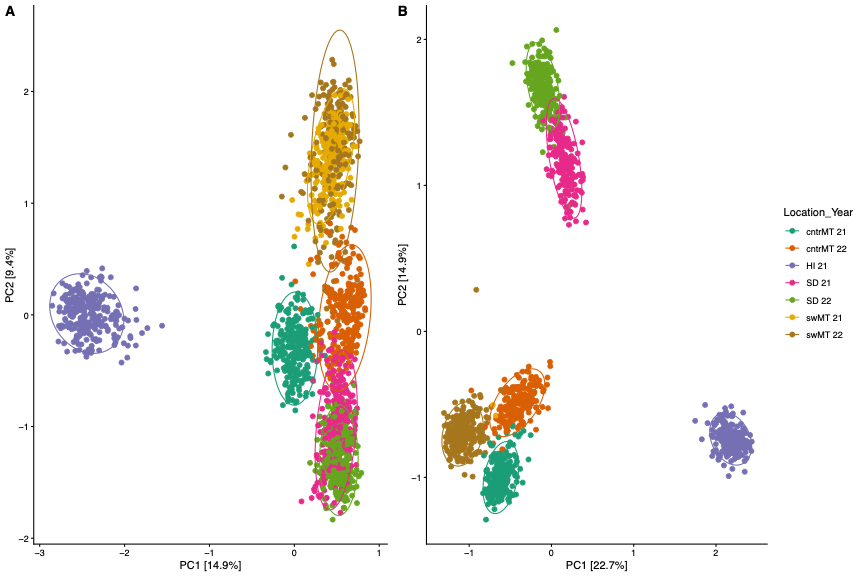

**Fig. S3.** Principal component analysis comparing bacterial (A) and fungal (B) communities between locations and years including Hawaii.

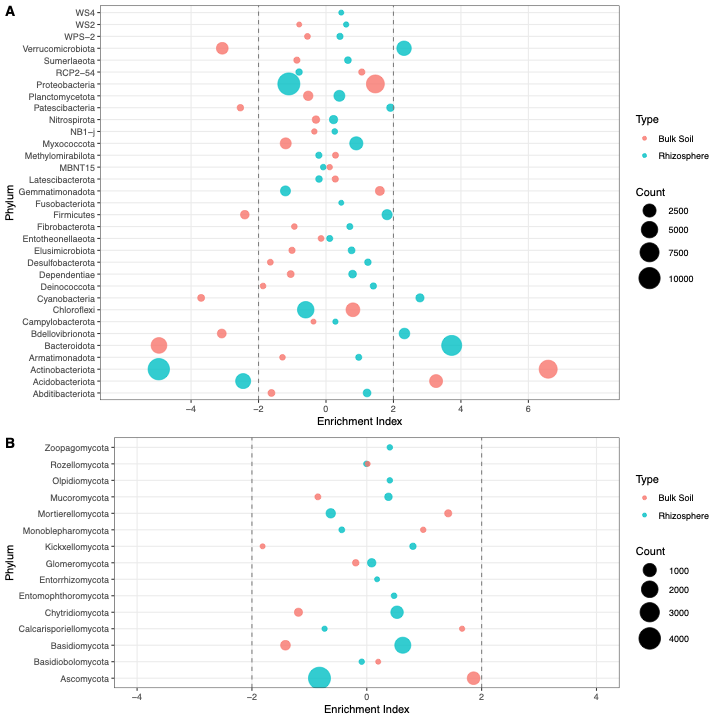

**Fig. S4.** Enrichment analysis at the phylum level comparing barley genotypes grouped by ecotype for rhizosphere bacterial (A) and fungal (B) communities Circle size indicates the number of ASVs representing a phylum for each cluster. Phyla with an enrichment index >2 or < -2 indicate significant over or underrepresentation within each cluster.

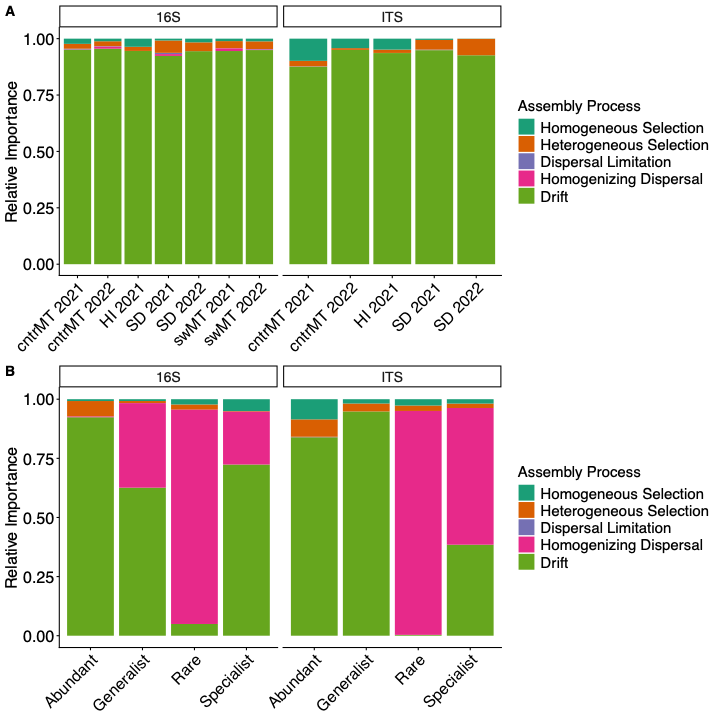

**Fig. S5.** Relative importance of stochastic and deterministic ecological processes in shaping rhizosphere bacterial and fungal community assembly compared among location-year (A) and ecotype (B).
